# Identifying the phenotypic effect of rare variants by including between-pedigree coancestry in variance components linkage analysis

**DOI:** 10.1101/215863

**Authors:** J.E. Hicks, M. A. Province

## Abstract

The contribution of rare variants to disease burden has become an important focus in genetic epidemiology. These effects are difficult to detect in population-based datasets, and as a result, interest in family-based study designs has resurfaced. Linkage analysis tools will need to be updated to accommodate the scale of data generated by modern genotyping and sequencing technologies.

In conventional linkage analysis individuals in different pedigrees are assumed to be independent of each other. However, cryptic relatedness is often present in populations and haplotypes that harbor rare variants may be shared between pedigrees as well as within them.

With millions of polymorphisms, Identity-by-descent (IBD) states across the genome can now be inferred without use of pedigree information. This is done by identifying long runs of identical-by-state genotypes which are unlikely to arise without IBD. Previously, IBD had to be estimated in pedigrees from recombination events in a sparse set of markers.

We present a method for variance-components linkage that can incorporate large number of markers and allows for between-pedigree relatedness. We replace the IBD matrix generated from pedigree-based analysis with one generated from a genotype-based method. All pedigrees in a dataset are considered jointly, allowing between-pedigree IBD to be included in the model.

In simulated data, we show that power is increased in the scenario when there is a haplotype shared IBD between members of different pedigrees. If there is no between-pedigree IBD, the analysis reduces to conventional variance-components analysis. By determining IBD states by long runs of dense IBS genotypes, linkage signals can be determined from their physical position, allowing more precise localization.

## Introduction

With the availability of high throughput genome sequencing, rare variants are an important new direction of investigation in genetic epidemiology. Detecting effects of rare variants is difficult in population-based study designs. Since individual rare variants account for little of the population heritability, family-based study designs may provide a more powerful to detect their effects. Population-based studies require extremely large sample sizes for a rare variant to be found in multiple participants. Due to their relatedness, a rare variant found in one pedigree is much more likely to be found in a relative than an individual from the general population.

Rare variants tend to be recent in origin, and lay on long haplotypes relatively undisturbed by recombination events. These regions are detectable in pairs of individuals directly from genotype data, and current tools for family-based genetics analysis will need to be updated to exploit this phenomenon.

Cryptic relatedness is a phenomenon in which individuals are related without their own knowledge. This is more common than expected from basic population genetic models (Gusev et al. 2012), but is consistent with population history (Henn et al. 2012; Palamara et al. 2012; Ralph and Coop 2013). In family-based analysis pedigrees are generally assumed to be independent from each other. However, pedigrees drawn from a single population can show cryptic relatedness as well. Multiple pedigrees can share a distant common ancestor. If an ‘unrelated’ pair of individuals share a segment of the genome containing a rare high-effect variant, this would be an important source of covariance that would go unmodeled in analysis methods that consider pedigrees separately.

Conventional methods of IBD determination cannot scale to dense genotypes in large pedigrees. The Lander-Green algorithm (Lander and Green 1987) enumerates all possible descent patterns to determine IBD status, limiting the size of pedigrees that can be analyzed. By working on each pedigree individually, it is limited to within-pedigree IBD, and cannot detect cryptic relatedness between pedigrees. Additionally, marker alleles must be in linkage equilibrium. To meet this requirement, the majority of genotype data available from SNP array or sequencing experiments must be discarded.

Kong et al. (2008) describe a general method of identifying IBD regions in a pair of individuals, in which IBD is determined by long consecutive stretches of genotypes shared Identical-by-state (IBS). These long IBS stretches are unlikely to occur in the absence of local common ancestry. A linkage model that incorporates IBD states determined in this manner would have several advantages. IBD determination in this fashion is less computationally complex, and extracts more precise pairwise inheritance information from the available genotypes than Lander-Green. Linkage regions can be localized to physical positions instead of chromosome locations in centimorgans. Finally, IBD can be identified between pedigrees. When between-pedigrees IBD is present, it is an important source of information that has otherwise been missed.

Here we propose a shared genomic segment method of variance components linkage (SGS-VC). IBD states are evaluated from runs of IBS, and all pedigrees in a dataset considered jointly. This model can be evaluated rapidly and shows increased power compared to conventional methods. We also show that when variants influencing a phenotype are shared between pedigrees, the inclusion of this covariance greatly increases power to detect linkage.

## Method

We consider the linear variance-components model for linkage analysis:

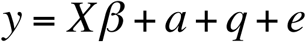

where X is a design matrix of fixed effects with corresponding coefficients β, a is the additive effect of polygenes, q is the effect of a QTL at the locus under consideration and e is residual error. Observations have the variance-covariance matrix:

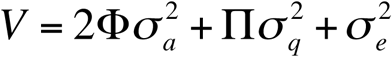

with Φ as the additive relationship matrix and Π is the matrix of IBD states at the locus. This model is fit by restricted maximum likelihood estimation. To determine statistical significance, the model is compared to a model lacking the QTL effect terms.

IBD states for the model can be determined by identifying shared genomic segments. While many implementations exist for this task (Browning and Browning 2011; Gusev et al. 2009; Han and Abney 2011; Hickey et al. 2011), we have chosen a simple method for determining IBD states. IBS states were evaluated for each genotype for every pair of genotyped individuals. Runs of IBS>0 longer than 1Mb were considered IBD=1 and runs of IBS=2 were considered IBD=2. This method does not require haplotype phase, avoiding substantial computation time and a potential source of error. Since the pedigree is not used to determine IBD, genotypes do not need to be assigned to ancestors, and this time consuming step is also avoided.

Without pedigree data, IBD can be determined for pairs of individuals across pedigrees, making computation time proportional to the number of pairs of genotyped individuals instead of pedigree size. When incorporating between-pedigree IBD, the number of pairs of individuals to evaluate is much greater and results in longer computation times.

This method evaluates all individuals in the dataset jointly. As a result the covariance matrices used in the mixed model can become large. Since fitting mixed models requires inverting these matrices, computation time for the statistical portion of this method is increased, but may result in increased power around potential causal variants.

### Simulated data

We evaluate the performance of this method using one hundred simulated datasets generated by the *pydigree* software package. Each replicate consisted of fifty pedigrees based on the template shown in Figure 1.

**Figure 1.**
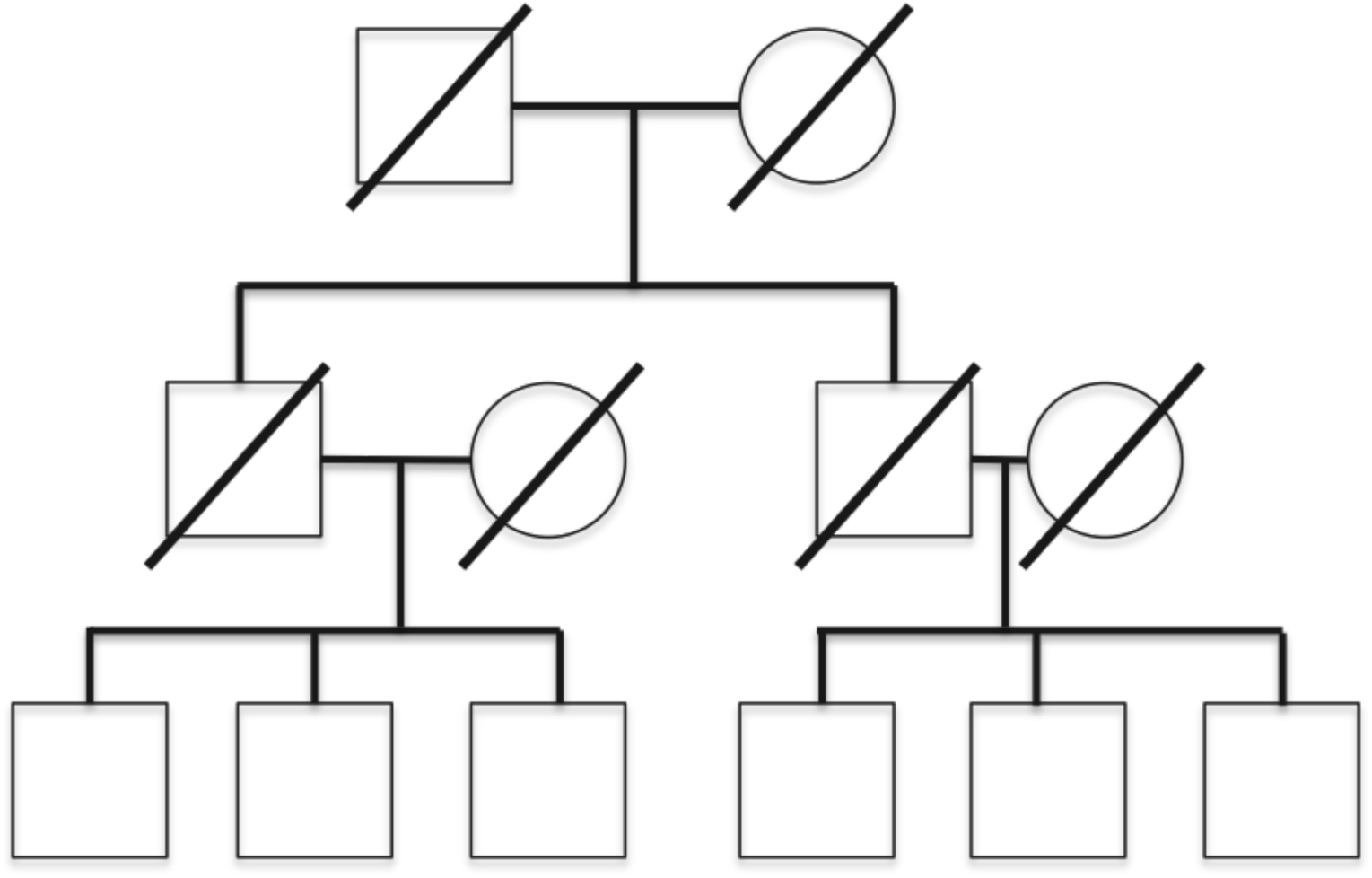
Pedigree used in simulations. Only the bottom generation had simulated genotypes and phenotypes.

The simulation phenotype was generated under a standard normal distribution (μ=0, σ=1), and was set to have a narrow-sense heritability of 50%. Genotypes were based on the SNPs available on chromosome 12 from the Affymetrix 6.0 platform. This resulted in a set of 30,882 common SNPs per individual. A SNP at 42,203,444 bp was designated as the causal variant. We create a 5,000 SNP haplotype that contained this variant. This haplotype was the only place the causative variant could be found. This procedure was for effect sizes of 0, 0.5, 1.0, 1.5, and 2.0.

Genotypes were simulated using a forward-time method. To include cryptic relatedness into the simulation, each pedigree in the simulation had a 20% chance of a founder individual carrying the causal haplotype. For these pedigrees, meiosis were constrained so that the causal haplotype was transmitted to at least one individual in each of the sibships.

### Comparison methodologies

We chose to compare the performance of SGS-VC to other methods. We consider two models: one including cryptic relatedness (SGS-VC-Joint+Between), and an identical model with between-pedigree IBD states fixed at 0 (SGS-VC-Joint). To compare with conventional linkage, analysis on the simulated dataset was performed using the variance-components model implemented in MERLIN (Abecasis et al. 2002). MERLIN assumes the set of marker genotypes is in linkage equilibrium. Therefore a subset of 133 SNPs were used for analysis.

The performance of association methods was determined by a variance-components association model, using the pedigree relationship matrix as a random effect and each SNP as a fixed effect. Since fitting the variance-components model is time-intensive, association was only evaluated at the causal variant. To account for the multiple testing burden of a genome-wide association strategy, the significance threshold was set at 10^−8^.

## Simulation Results

In all simulations, type I error was well controlled for each method. Even in the presence of between pedigree IBD, there was no increase in false positive rate when no causal variant was present.

Power to determine evidence of linkage at LOD score thresholds of 2 and 3 are given in tables 1 and 2, respectively. In the presence of a high-effect variant SGS-VC had substantially more power across a range of effect sizes to detect linkage than MERLIN or an SGS-VC-Joint. MERLIN showed low power for every effect size. As effect sizes increased, SGS-VC-Joint showed increased power compared to MERLIN. For a LOD threshold of 3, SGS-VC-Joint+Between was the only model that showed good power for effect sizes less than 2. Mixed model association showed good power to detect the variant at high effect sizes, but was lower than the SGS-VC-Joint+Between.

**Table 1.** Power of methods in simulation data to reach a LOD score greater than 2.

| Effect Size | MERLIN | SGS-VC-Joint | SGS-VC-Joint+Between |
| --- | --- | --- | --- |
| 0.5 | 0.01 | 0 | 0.08 |
| 1.0 | 0.04 | 0.06 | 0.69 |
| 1.5 | 0.16 | 0.32 | 0.98 |
| 2.0 | 0.61 | 0.82 | 1.0 |

**Table 2.** Power of methods in simulation data to reach genome-wide significance (LOD > 3 for linkage methods, *p* < 10^−8^ for association).

| Effect Size | MERLIN | SGS-VC-Joint | SGS-VC-Joint+Between | Mixed model association |
| --- | --- | --- | --- | --- |
| 0.5 | 0 | 0 | 0.03 | 0 |
| 1.0 | 0 | 0 | 0.53 | 0.23 |
| 1.5 | 0.05 | 0.11 | 0.95 | 0.89 |
| 2.0 | 0.38 | 0.65 | 1.0 | 1.0 |

## Discussion

In general, the use of IBD states determined by SGS outperformed MERLIN. In the presence of between pedigree IBD, SGS-VC-Joint+Between showed much higher power to detect linkage than other methods, indicating that modelling between-pedigree covariance can be an important factor in identifying causative alleles.

Even without incorporating between-pedigree IBD, SGS-VC-Joint remains a powerful method of linkage. Consideration of the dataset as a whole shows higher evidence of linkage, even when only a fraction of pedigrees in the dataset have the causative allele. When between-pedigree IBD is ignored, computation can be completed rapidly, as the number of pairs to evaluate is much smaller. If covariance due between-pedigree IBD is incorporated but not present, SGS-VC-Joint+Between reduces to the SGS-VC-Joint model.

In simulated data, mixed-model association had similar power to SGS linkage. However, this requires testing of the causal variant directly. Fitting the variance-component model requires inverting large matrices, a computationally intensive process that makes it an infeasible to test rare variants at genome-wide scale. Instead, association should be a complementary part of an analysis. As a linkage method, SGS-VC performs well even when performed at a limited number of sites in the genome. Once a linkage signal is detected, association methods can be performed to further narrow the linkage region and identify individual variants.

This method shares some similarity to a method proposed by Day-Williams et al. (2011), but there are important differences. This method uses a linear model, which can easily be extended into more complicated models. We use pedigree structures as given, instead of re-estimating them, avoiding a potential loss of power. Finally, we deterministically generate IBD states, without of estimating using allele frequencies, avoiding another avenue of model misspecification.

Between-pedigree relatedness can be a useful source of information when trying to identify causative loci. As the number of pedigrees in a dataset increases, so does the likelihood of cryptic relatedness. SGS-VC presents a powerful way of exploiting that cryptic relatedness.

## Works Cited

Abecasis GR, Cherny SS, Cookson WO, Cardon LR (2002) Merlin--rapid analysis of dense genetic maps using sparse gene flow trees. Nat Genet 30: 97–101. doi: 10.1038/ng786

Almasy L, Blangero J (1998) Multipoint quantitative-trait linkage analysis in general pedigrees. Am J Hum Genet 62: 1198–211. doi: 10.1086/301844

Amos CI (1994) Robust variance-components approach for assessing genetic linkage in pedigrees. Am J Hum Genet 54: 535–43.

Browning BL, Browning SR (2011) A fast, powerful method for detecting identity by descent. Am J Hum Genet 88: 173–82. doi: 10.1016/j.ajhg.2011.01.010

Day-Williams AG, Blangero J, Dyer TD, Lange K, Sobel EM (2011) Linkage analysis without defined pedigrees. Genet Epidemiol 35: 360–70. doi: 10.1002/gepi.20584

Gusev A, Lowe JK, Stoffel M, Daly MJ, Altshuler D, Breslow JL, Friedman JM, Pe’er I (2009) Whole population, genome-wide mapping of hidden relatedness. Genome Res 19: 318–26. doi: 10.1101/gr.081398.108

Gusev A, Palamara PF, Aponte G, Zhuang Z, Darvasi A, Gregersen P, Pe’er I (2012) The architecture of long-range haplotypes shared within and across populations. Mol Biol Evol 29: 473–86. doi: 10.1093/molbev/msr133

Han L, Abney M (2011) Identity by descent estimation with dense genome-wide genotype data. Genet Epidemiol 35: 557–67. doi: 10.1002/gepi.20606

Henn BM, Hon L, Macpherson JM, Eriksson N, Saxonov S, Pe’er I, Mountain JL (2012) Cryptic distant relatives are common in both isolated and cosmopolitan genetic samples. PLoS One 7: e34267. doi: 10.1371/journal.pone.0034267

Hickey JM, Kinghorn BP, Tier B, Wilson JF, Dunstan N, van der Werf JH (2011) A combined long-range phasing and long haplotype imputation method to impute phase for SNP genotypes. Genet Sel Evol 43: 12. doi: 10.1186/1297-9686-43-12

Kong A, Masson G, Frigge ML, Gylfason A, Zusmanovich P, Thorleifsson G, Olason PI, Ingason A, Steinberg S, Rafnar T, Sulem P, Mouy M, Jonsson F, Thorsteinsdottir U, Gudbjartsson DF, Stefansson H, Stefansson K (2008) Detection of sharing by descent, long-range phasing and haplotype imputation. Nat Genet 40: 1068–75. doi: 10.1038/ng.216

Lander ES, Green P (1987) Construction of multilocus genetic linkage maps in humans. Proc Natl Acad Sci U S A 84: 2363–7.

Palamara PF, Lencz T, Darvasi A, Pe’er I (2012) Length distributions of identity by descent reveal fine-scale demographic history. Am J Hum Genet 91: 809–22. doi: 10.1016/j.ajhg.2012.08.030

Ralph P, Coop G (2013) The geography of recent genetic ancestry across Europe. PLoS Biol 11: e1001555. doi: 10.1371/journal.pbio.1001555

